# Organic farming expansion drives natural enemy abundance not diversity in agricultural landscapes

**DOI:** 10.1101/688481

**Authors:** Lucile Muneret, Arthur Auriol, Olivier Bonnard, Sylvie Richart-Cervera, Denis Thiéry, Adrien Rusch

## Abstract

1. Organic farming is seen as a prototype of ecological intensification able to conciliate crop productivity and biodiversity conservation in agricultural landscapes. However, how natural enemies, an important functional group supporting pest control services, respond to organic farming at different scales and in different landscape contexts remain unclear.
2. Using a hierarchical design within a vineyard-dominated region located in southwestern France, we examine the independent effects of organic farming and semi-natural habitats at the local and landscape scales on natural enemies.
3. We show that the proportion of organic farming is a stronger driver of species abundance than the proportion of semi-natural habitats and is an important facet of landscape heterogeneity shaping natural enemy assemblages. Although our study highlight a strong taxonomic group-dependency about the effect of organic farming, organic farming benefits to dominant species while rare species occur at the same frequency in the two farming systems.
4. Independently of farming systems, enhancing field age, reducing crop productivity, soil tillage intensity and pesticide use are key management options to increase natural enemy biodiversity.
5. *Synthesis and Applications*. Our study indicates that policies promoting the expansion of organic farming will benefit more to ecological intensification strategies seeking to enhance ecosystem services than to biodiversity conservation.

## 1 Introduction

Biodiversity and agricultural production play a vital role in human society (Cardinale et al., 2012). However, a huge trade-off exists between these two inextricably linked components. Their relationships have been thought in regard to the land sharing - land sparing debate that aims at identifying the best strategies for conciliating them (Phalan, Onial, Balmford & Green, 2011). To date, no clear consensus exists on the best landscape planning strategy to optimize synergies between crop production and biodiversity conservation. Two reasons among others are that it is highly taxon-dependent (Martin, Seo, Park, Reineking & Steffan-Dewenter, 2016) and depends on the spatial scale on which it applies (Ekroos et al., 2016). In addition to conservation goals, biodiversity is generally associated with greater levels of ecosystem functioning and thus offers a path to the development of ecological intensification of agricultural systems (Cardinale et al., 2012; Bommarco, Kleijn & Potts, 2013).

Although no ideal model of ecological intensification does exist, organic farming is often seen as a good prototype (Tittonnell, 2014). While organic farming is on average 19-25% less productive than conventional fields (Ponisio et al., 2015), it supports higher levels of biodiversity and ecosystem services than conventional farming (Muneret et al., 2018a, Tuck et al., 2014). Nonetheless, the effect of organic farming on biodiversity and ecosystem services is found to be highly context dependent but we lack a good understanding about the mechanisms underlying this context dependency (Tuck et al., 2014; Tscharntke et al., 2012).

One hypothesis beyond the context-dependency of organic farming performances on biodiversity and ecosystem services is the potential interactions between local farming practices and the surrounding landscape (Tscharntke et al., 2012; Kleijn, Rundlöf, Scheper, Smith & Tscharntke, 2011). To date, studies examining how landscape context modulates the local effect of farming practices for biodiversity conservation or the provision of ecosystem services have mainly considered the impact of semi-natural habitats but much less attention have been paid to the role of farming practices within the landscape (Rusch et al., 2016; Chaplin-Kramer, O’Rourke, Blitzer & Kremen, 2011; but see Henckel, Börger, Meiss, Gaba & Bretagnolle, 2015). Moreover, a recent synthesis at the global scale revealed strong variability in direction and effect size of semi-natural habitats on predators and biological pest control (Karp et al., 2018). Thus, considering farming practices at multiple spatial scales, notably the proportion of organic farming at the landscape scale, should reduce unexplained variation in these relationships (Muneret, Auriol, Thiéry & Rusch, 2019).

Studies on the effect of organic farming have mostly been conducted at the field scale and the few studies conducted at a landscape scale highlighted that the response of taxon is highly idiosyncratic (Gabriel et al., 2010). Moreover, the effects of semi-natural habitats and organic farming in the landscape could interact because they could potentially support different set of species having either antagonistic, synergistic or even neutral interactions (Letourneau, Jedlicka, Bothwell & Moreno, 2009; Martin, Reineking, Seo & Steffan-Dewenter, 2013). Hence, examining the effect of organic farming applied at much larger scales is urgently needed since organic agriculture is expanding exponentially and receives important subsidies, notably in Europe, while very limited knowledge about the performance of such cropping systems in the context of large spatial expansion (Kleijn et al., 2011).

Based on a sampling design composed by 42 commercial vineyards, we independently investigated the effects of organic farming and semi-natural habitats at multiple scales on natural enemy biodiversity. The few studies examining the effect of the proportion organic farming on natural enemies never disentangled the effects of organic farming from the effect of semi-natural habitats at the landscape scale while evidence shows that they are correlated (Norton et al., 2009). Moreover, perennial crops are far less studied than annual ones while their own cropland biodiversity could be differentially affected by agricultural operations (Bruggisser, Schmidt-Entling, Bacher, 2010). Such crops provide more refugees and resources for biodiversity but generally received many more pesticide applications over time than annuals crops (Muneret, Thiéry, Joubard & Rusch, 2018b). Here, we sampled a wide community of arthropod natural enemies (spiders, harvestmen, ground beetles, rove beetles, lacewings, ants and earwigs). First, we aim at disentangling and evaluating the relative effect of the ‘hidden heterogeneity’ (here referred as the proportion of organic farming; Vasseur et al., 2013) from the relative effect of the proportion of semi-natural habitats on the natural enemy community. Second, we examine the effect of interaction between local farming systems and the landscape context on the arthropod biodiversity. Third, we identified several farming practices related to the local management intensity that impact arthropod biodiversity. Finally, we discuss how the expansion of organic farming, as a land-sharing path, can outline trade-offs between conservation goals and ecological intensification application in agricultural landscapes.

## 2 MATERIAL AND METHODS

### 2.1 Study design

Our study design was located within a vineyard-dominated region, in South-Western France (near Bordeaux, 44°81’N, −0°14’W). The design consisted 42 vineyard plots organized in 21 pairs, each pair containing one field managed under organic guidelines and one not (hereafter referred as “conventional”). The average distance between paired fields was about 125 m. Pairs of vineyards have been selected along two uncorrelated landscape gradients: proportion of semi-natural habitats and proportion of organic farming. These gradients were established based on landscape composition calculated at a 1 km radius around each focal vineyard. At this scale, the proportion of semi-natural habitats ranged from one to 75%, and the proportion of organic farming ranged from two to 25% of the total land area. This study design allowed for the unraveling of farming system effects at the local scale as well as the relative effects of the proportions of semi-natural habitats and organic farming on biodiversity. Landscape variables were also calculated at the 500-m radius around each vineyard using ArcGIS 10.1 (ESRI).

### 2.2 Arthropod sampling

We sampled natural enemy communities on the soil surface and in the foliage. The community of the soil surface was sampled by placing on the ground five pitfall traps per vineyard (diameter 11 cm; depth 11.5 cm) under three vine rows that were distant from three vine rows from each other (inter-row distances varied between 1.5 and 3 m). Three pitfall traps were placed at 10-m and two others at 10-m away from the edge. They were opened during seven days at five sampling dates between late May and early October in 2015. The five traps were pooled at each sampling period prior to analyses. In addition, the foliage community was sampled four times between early June and early September by beating 30 vine stocks at least 5 vine stocks away from each other along two or four vine rows depending on the field size. Harvestmen, spiders, ants, rove beetles and ground beetles were identified at the species level while lacewings and earwigs were identified at the family level. Ant community collected in pitfall traps were only counted and identified at the first sampling date (i.e. in early June).

### 2.3 Vineyard management

We collected data about pesticide use (i.e., fungicide, insecticide and herbicide), soil tillage and field age by interviewing the 38 vine growers involved. The intensity of pesticide application was calculated using the Treatment Frequency Index which is the sum of all the ratios between the applied and the recommended dose for each pesticide application (OECD, 2001). Tillage intensity was evaluated by calculating the “tillage intensity index” which summarizes the number of tilling operations per year weighted by the area involved each time (see Muneret et al., 2019). We also measured vine trunk density and crop productivity. To calculate the crop productivity, we multiplied the average number of bunches per vine stock by the average bunch weights and the vine stock density per vineyard (Mg/ha, see Muneret et al., 2018b). Note that crop productivity did not significantly differ between farming systems and that we were not able to estimate crop productivity for two vineyards out of 42 (see Muneret et al., 2018b).

### 2.4 Data analyses

As the above-ground community and the foliage community represent two guilds, we analysed the response of each community to environmental conditions separately. For each community, we calculated total abundance, species richness and evenness (Pielou index) over the year. At each sampling date, approximately 10% of the vineyards were not sampled because of pitfall trap destruction (N=4 for the first until the fourth sampling dates and N=5 for the fifth date). Species richness was rarefied to take into account differences in terms of detectability within fields (Gotelli & Colwell, 2001). For the foliage community, we calculated the abundance of ants, spiders, earwigs and lacewings and the richness of ants and spiders. For the above-ground community, we calculated both the abundance and the richness of ants, spiders, ground beetles and rove beetles. Finally, we calculated the global abundance of harvestmen (those from both the foliage and the ground). All these metrics represented 21 descriptors of natural enemy communities which were then used as response variables in our models and data were log-transformed for further analyses when it was necessary.

Linear mixed models were used to investigate the effects of local management intensity, farming systems and landscape composition on each response variable. Because of some trap destructions, we corrected the abundance of the communities for each vineyard having an uncompleted sampling. Therefore, separately for the foliage and the above-ground community, we calculated the relative contribution of each sampling date to the total abundance of the community and we divided the total number of individuals collected by the sum of the relative contribution of the sampling dates that were sampled for the given vineyard. This gave the estimated total abundance of a given community for a given vineyard taking account for which sampling dates were sampled.

We fitted four models of increasing complexity (‘M0’, ‘M1’, ‘M2’ at 500-m scale and ‘M2’ at the 1000-m scale) for each response variable and we used a multimodel inference approach to test our hypotheses (N_*observations*_=40). We applied this procedure to identify the most relevant spatial scale for natural enemies. M0, the first model, had local covariates as predictors: ‘field age’, ‘vine stock density’, ‘total TFI’, ‘tillage intensity’ and ‘crop productivity’. At this first step, all the possible models were ranked using the Akaike Information Criteria corrected for small sample size (AICc) and models with a ΔAICc < 2 were retained among the set of top models. Such set of top models was then used to estimate the mean effects and confidence intervals of each explanatory variable using model averaging (Grueber, Nakagawa, Laws & Jamieson, 2011). Covariates which were significant at this M0 step (*i.e.*, with a confidence interval significantly different from zero and having a relative importance variable equal to 1) were conserved and included in models ‘M1’. M1 included ‘selected local covariates’ and ‘local farming systems’ as predictors. This step allows for evaluating the effect of local farming systems on biodiversity after taking into account potential confounding effects of specific local covariates. We then fitted two different M2 models, one for each spatial scale (*i.e.*, 500-m, and 1,000-m) to test our hypotheses related to the effect of the landscape composition and its interaction with local farming systems on biodiversity. In M2, we thus integrated ‘selected covariates’, ‘local farming systems’, ‘the proportion of semi-natural habitats’, ‘the proportion of organic farming’ and two interactions: (i) local farming systems with the proportion of semi-natural habitats, (ii) local farming systems with the proportion of organic farming. All the models at all scales included ‘field pairs’ as random effect.

The same averaging approach was applied for the four models and we calculated the marginal R^2^ values and conditional R^2^ values of the model having the lowest AICc at each step to evaluate the amount of variability explained by each of the six models (Nakagawa & Schielzeth, 2013). Before modeling, we standardized all explanatory variables, with mean equal to 0 and standard deviation equal to 0.5 (Schielzeth, 2010).

To identify which level of model complexity, and indirectly which spatial scale, was the most important for explaining natural enemy descriptors, we recalculated the Akaike weights (‘Sum Wi’) among all of the models from the four different sets (*i.e.*, M0, M1 and M2 at both spatial scales) obtained for each response variable. We therefore estimated the relative importance of each level of complexity for a given response variable. The sum of the Akaike weights of the models obtained at a given level of complexity provided the model’s probability of being top model across all scales.

Diagnostic residual plots of all full models were confirmed using the DHARMa package (Hartig, 2017). Using variograms, we detected no spatial autocorrelation in the residuals. Collinearity among explanatory variables was assessed using the Variance Inflation Factor and the highest value was equal to 2.29 for the TFI.

All analyses were performed using the R software (R Core Team, 2016) and the packages ‘lme4’ (Bates, Mächler, Bolker & Walker, 2014) and ‘MuMIn’ (Bartoń, 2016).

## 3 RESULTS

### 3.1 Description of the natural enemy communities

We identified 41,663 arthropods belonging to 318 taxa. We collected 15,316 spider adults and juveniles (162 taxa), 5,074 ground beetle adults (60 taxa), 1,574 rove beetle adults (47 taxa), 16,911 ant adults (41 taxa), 1,864 harvestman adults and juveniles (6 taxa), 650 earwig adults (one family) and 274 lacewing larvae (one family). 19,549 individuals were collected in the foliage and 22,114 individuals were collected on the soil surface. Across all the 21 descriptors of the natural enemy community, models ‘M0’ with the simplest level of complexity (i.e. including local covariates only) mostly had the highest relative importance of explaining species richness of natural enemy community while their abundances mostly responded to models ‘M2’ with the highest level of complexity (i.e. including local covariates, local farming systems, landscape variables and interactions; Tables 1 et 2; Figure S1).

**TABLE 1.**
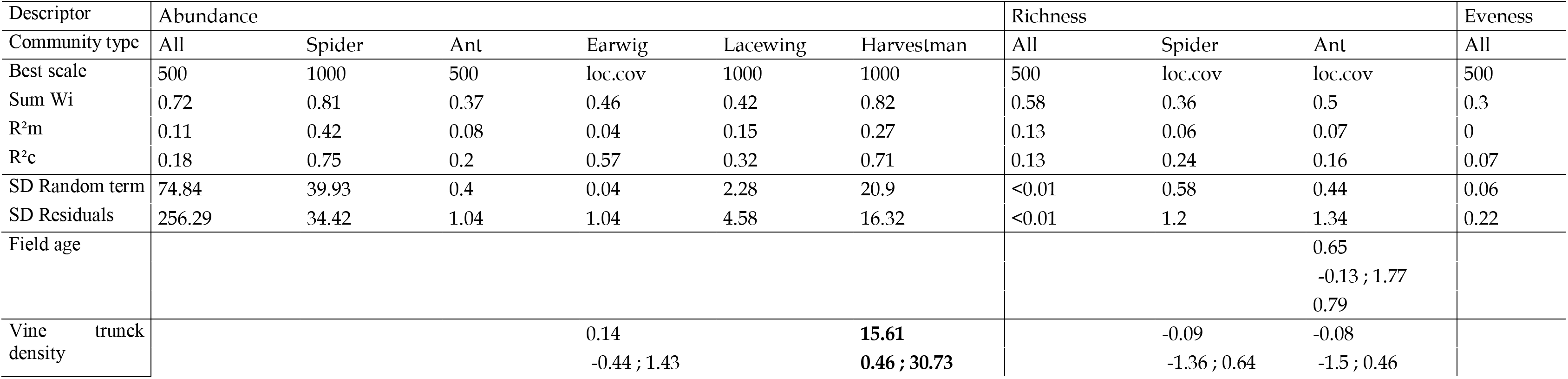
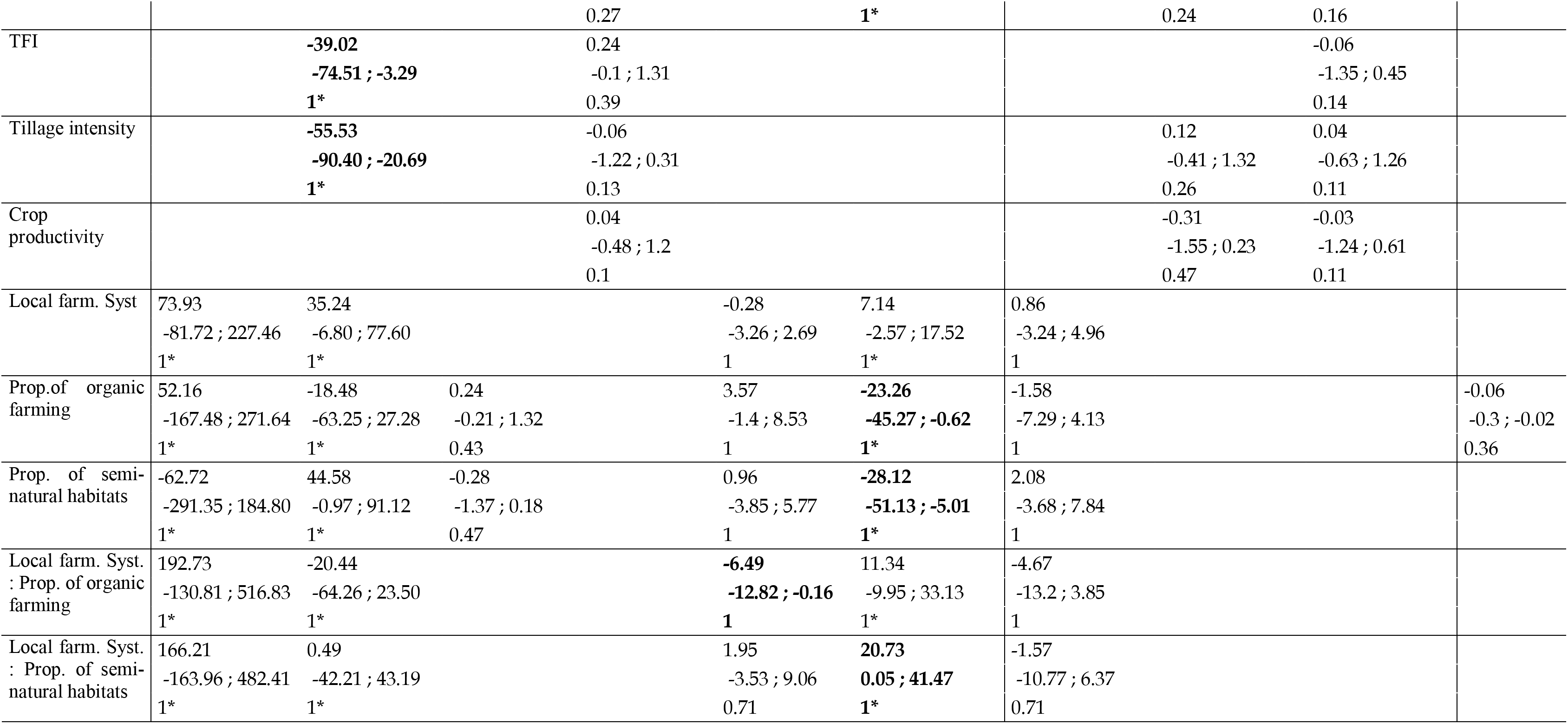
Responses of the natural enemy abundance, richness and evenness to environmental conditions in vineyards for the community sampled in the foliage. For each response variable, we recalculated the Akaike weights among all of the models from the four different sets (‘M0’, ‘M1’ and both ‘M2’ obtained using multimodel inference). We, therefore, estimated the relative importance of each level of complexity for a given response variable that gave us the “best scale” of response (Figure S1). The sum of the Akaike weights (“Sum Wi”) of the models obtained at the best scale provided the model’s probability of being the top model across all of the scales. Other parameters reported in this table come from models of the best scale for each response variable. R^2^ marginal and R^2^ conditional are reported. R2 values were calculated using the best models at the best scale. The standard deviation of the random terms are reported. Estimates, confident interval (2,5 - 97.5%) and relative importance variable were reported for each predictor. Values in bold are significant (confident interval did not include zero and relative variable importance equal to 1).

**TABLE 2.**
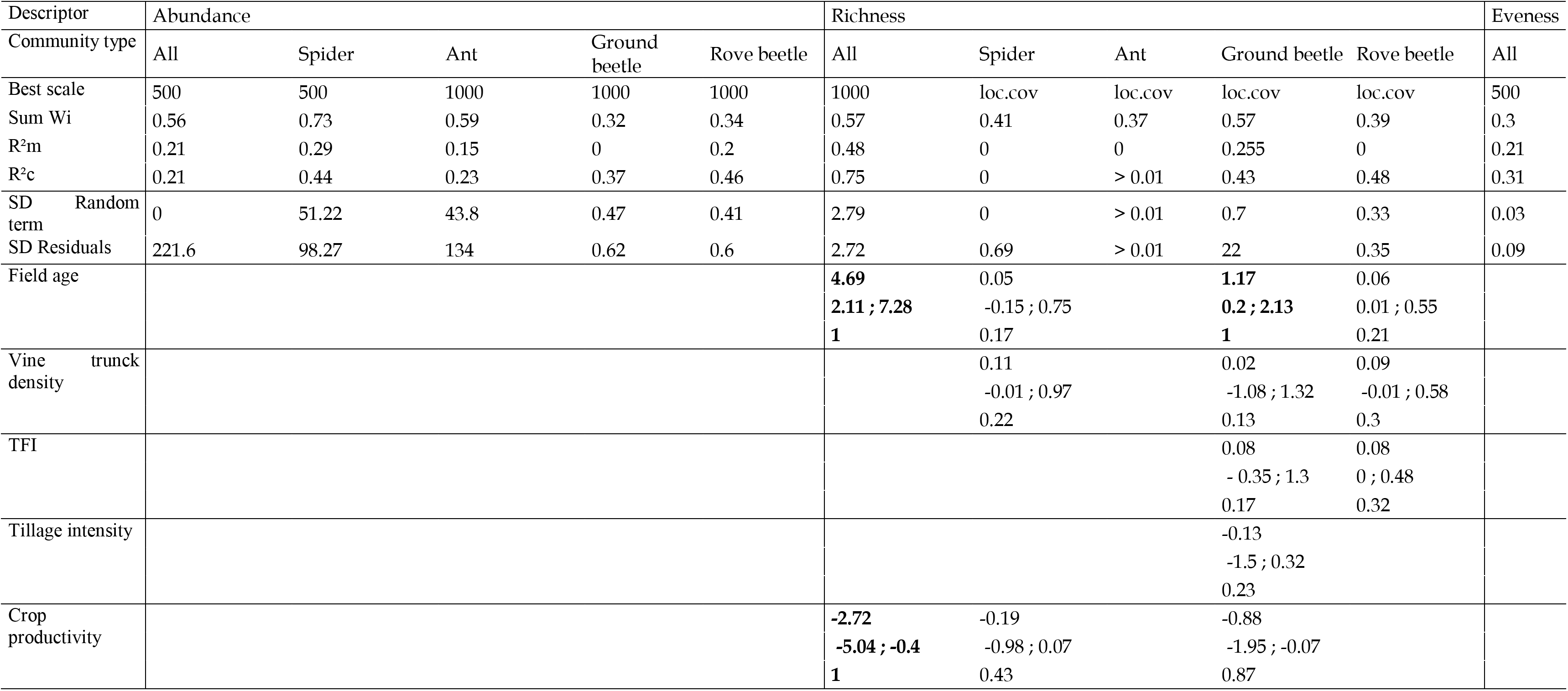
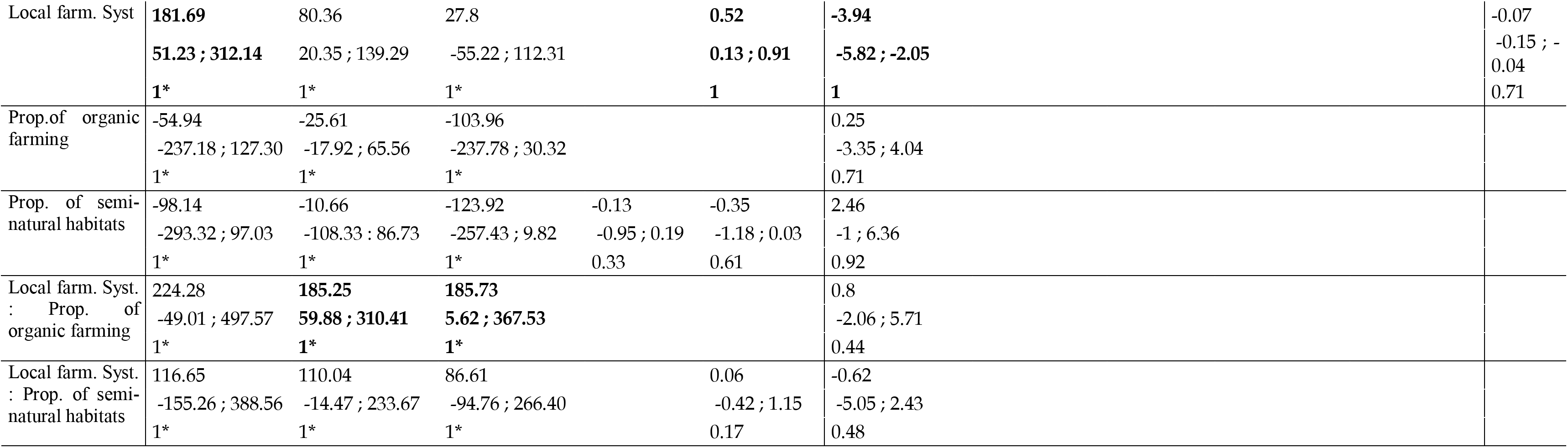
Responses of the natural enemy abundance, richness and evenness to environmental conditions in vineyards for the community sampled at the soil surface. All model descriptors that are reported have been obtained using the same procedure as the data reported in the Table 1. See the legend in Table 1.

### 3.2 Independent effects of local farming systems and landscape composition on natural enemy communities

At the field scale, organic farming did not impact the foliage community whereas it affected the above-ground community in several ways. Local organic farming increased the abundance of the total above-ground natural enemy community, as well as spider and rove beetle abundances (Table 2; Figure 1a). However, local organic farming negatively affected the total richness of the above-ground community (Table 2; Figure 1b). The three most abundant species (*Pardosa proxima*, *Lasisus niger* and *Pseudoophonus rufipes*) are much more abundant in organic than in conventional vineyards (Figure S2) what decrease the evenness of the above-ground community (Table 2, Figure 1c). The rarest species (i.e. species that were collected once) appear at the same frequency in organic and conventional vineyards (67 and 66 species in organic and conventional vineyards).

**FIGURE 1.**
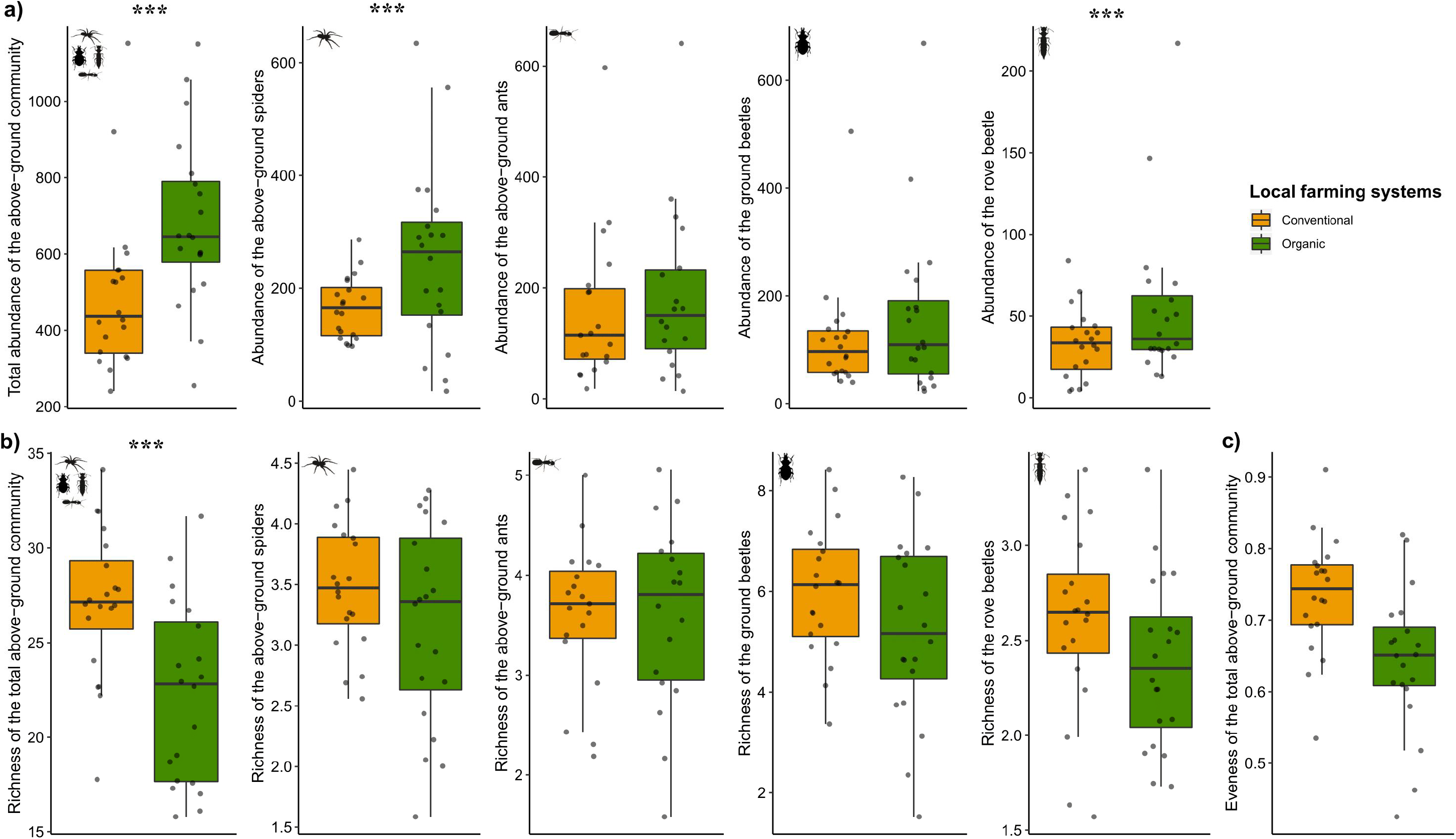
Effect of local farming systems on a) abundance, b) richness and evenness of the above-ground natural enemy community.

At the landscape scale, the proportion of semi-natural habitats and the proportion of organic farming, both alone, were negatively correlated with the abundance of harvestmen (M2 at the 1000-m scale had Sum Wi = 0.82; Relative variable importance = 1; R^2^_m_=0.27 and R^2^_C_=0.71; Figures 2a – 2b).

**FIGURE 2.**
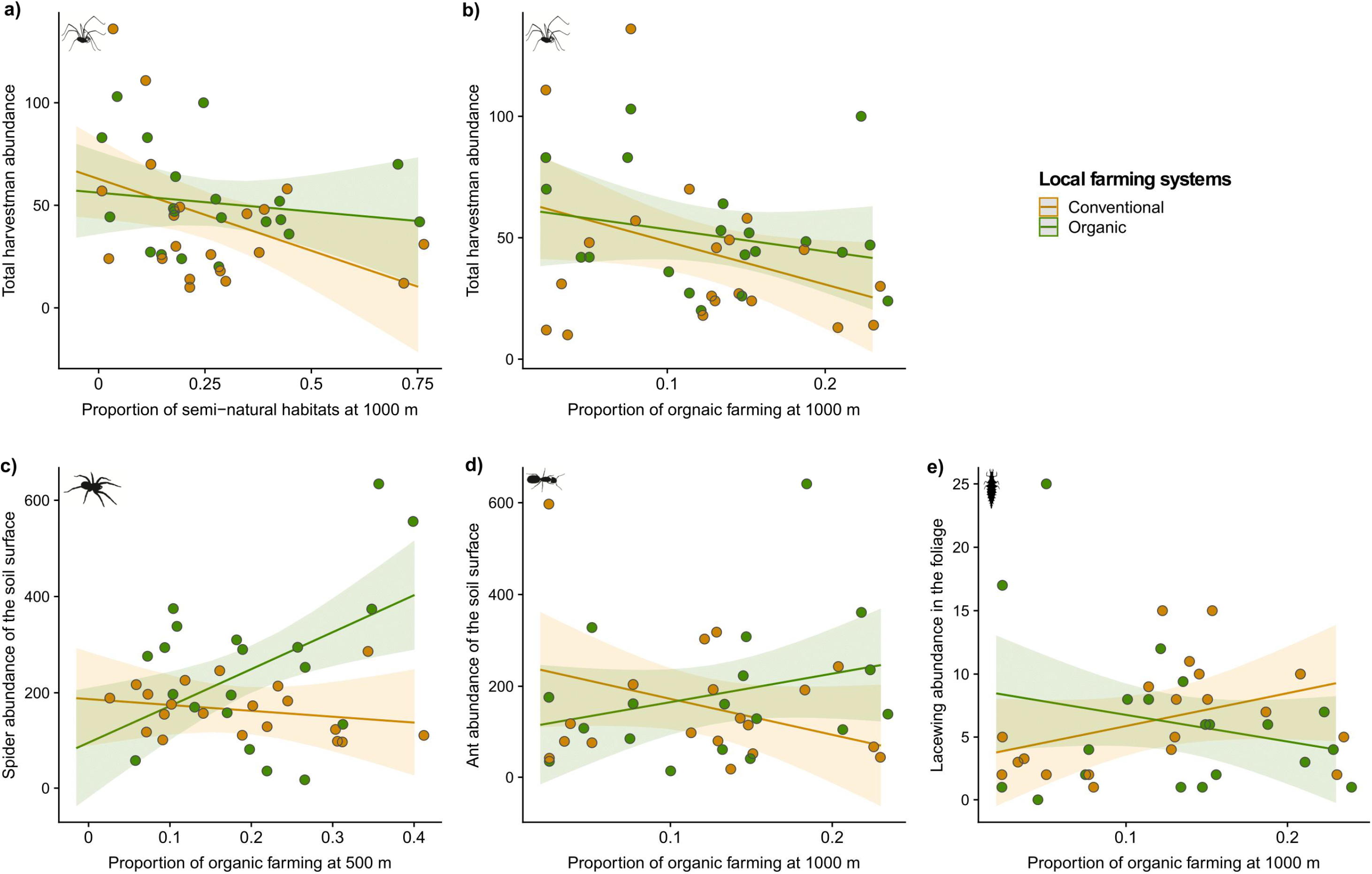
Interactive effects of landscape composition and local farming systems on natural enemy abundances. a) effect of the proportion of semi-natural habitats at the 1000-m scale and local farming systems on harvestman abundance; b) effect of the proportion of organic farming at the 1000-m scale and local farming systems on harvestman abundance; c) effect of the proportion of organic farming at the 500-m scale and local farming systems on above-ground spider abundance; c) effect of the proportion of organic farming at the 100-m scale and local farming systems on d) above-ground ant abundance and e) lacewing abundances.

### 3.3 Interactive effects of local organic farming and landscape composition on natural enemy communities

Three response variables responded to interactions between local farming systems and the proportion of organic farming (i.e. abundances of spiders and ants of the above-ground community and lacewing abundance) and one response variable responded to interactions between local farming systems and the proportion of semi-natural habitats (i.e. harvestman abundance). Contrary to abundance, natural enemy richness and evenness were not affected by interactions between local farming systems and landscape composition.

Abundances of spiders and ants of the above-ground community benefited more from local organic farming in landscapes having a higher proportion of organic vineyards in the landscape. For spiders, M2 at the 500-m scale had a Sum Wi = 0.73, a relative variable importance = 1, R^2^_m_=0.29 and R^2^_C_=0.44 (Table 2; Figure 2c). For ants, M2 at the 1000-m scale had a Sum Wi = 0.59, a relative variable importance = 1, R^2^_m_=0.15 and R^2^_C_=0.23 (Table 2; Figure 2d). On the opposite, lacewing abundance benefited more from conventional farming in landscapes having a high proportion of organic vineyards in the landscape (M2 at the 1000-m scale had a Sum Wi = 0.42, a relative variable importance = 1, R^2^_m_=0.15 and R^2^_C_=0.32; Table 1; Figure 2e).

Finally, harvestman abundance benefited more from organic farming in landscape having a high proportion of semi-natural habitats than in simpler landscapes ((M2 at the 1000-m scale had Sum Wi = 0.82; Relative variable importance = 1; R^2^_m_=0.27 and R^2^_C_=0.71; Table 1; Figure 2a).

### 3.4 Effect of the local management intensity on natural enemy communities

Independently of local farming systems, specific farming practices were important predictors of natural enemy community structure. We found positive effects of field age and vine trunk density on natural enemy communities. Field age had a positive effect on the richness of the total above-ground community and on the ground beetle richness (Figures 3a – 3b; Table 2) and that vine trunk density was positively correlated with harvestman abundance (Figure 3c, Table 1). On the opposite, crop productivity, tillage intensity and pesticide use intensity were negatively affected natural enemy communities. Crop productivity reduced the richness of the total above-ground community (Figure 3d, Table 2) while tillage intensity and pesticide use intensity had a negative impact on the abundance of spiders of the foliage (Figures 3e – 3f, Table 1).

**FIGURE 3.**
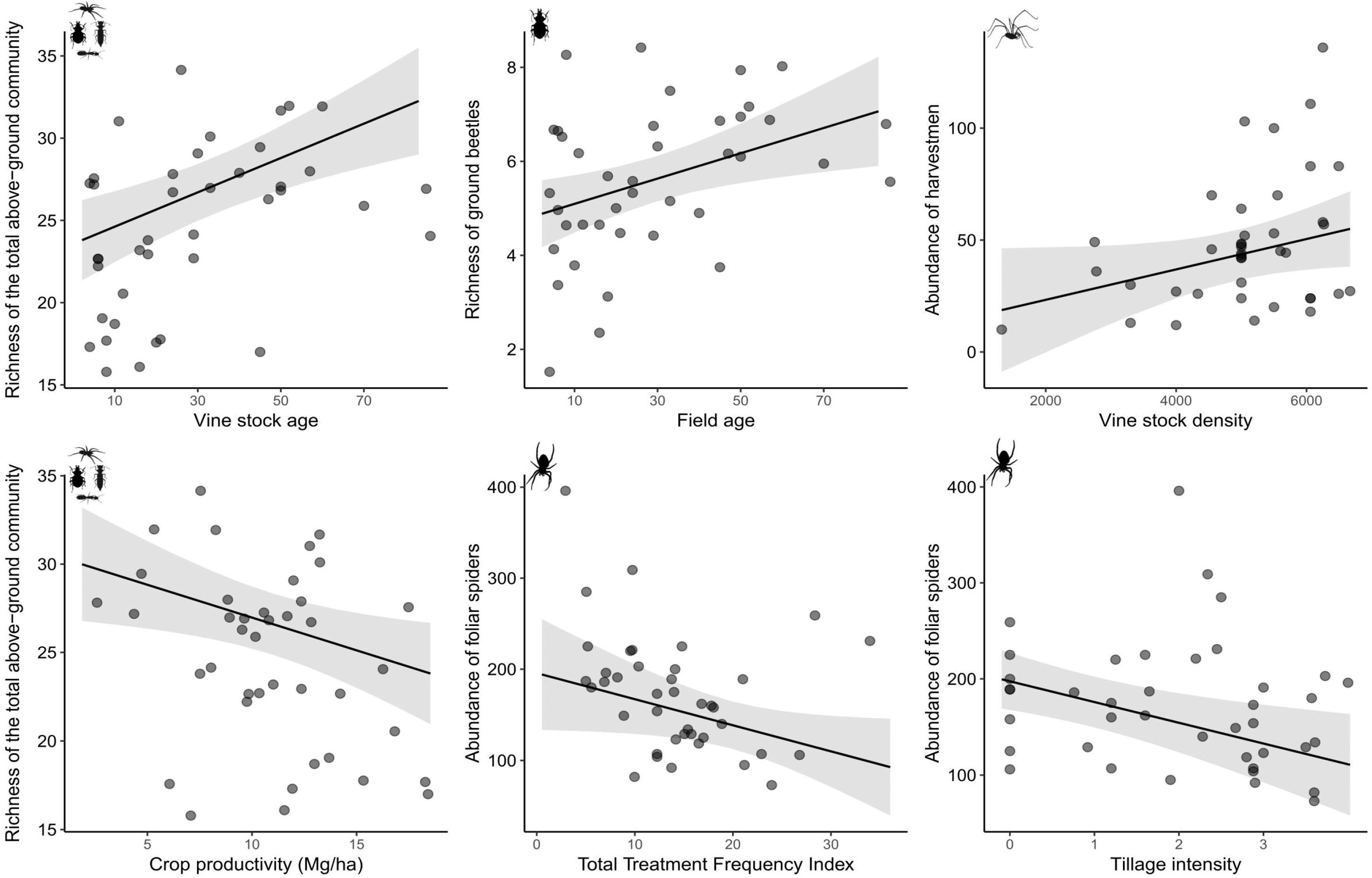
Effect of local covariates on abundance and richness on natural enemies. Effect of field age on a) the rarefied richness of the total above-ground community and b) the rarefied richness of the ground beetles. c) effect of vine trunk density on harvestman abundance. d) effect of crop productivity on the rarefied richness of the total above-ground community. Effect of e) treatment frequency index and f) tillage intensity on the abundance of the spiders of the foliage.

## 4 DISCUSSION

This study provides novel results about the independent effects of organic farming and semi-natural habitats at different spatial scales on natural enemy communities, a key functional group for the development of ecological intensification. Globally, we find that farming practices at multiple spatial scales are stronger drivers of community abundance than the proportion of semi-natural habitats. However, there is no unique response of arthropod communities to land use changes in agricultural landscapes. Most dominant agrobiontes species are more abundant in organic than in conventional farming while occurrence of the rarest species is the same between farming systems. Moreover, independently of farming systems, specific practices such as enhancing field age, reducing crop productivity, soil tillage intensity and pesticide use are key management options to increase natural enemy biodiversity. Our results suggest that organic farming which clearly enhances abundances of dominant predator species supports ecological intensification strategies but might not be sufficient enough to support arthropod conservation.

Farming systems at multiple spatial scales are a key determinant of natural enemy community structure in vineyard landscapes. This suggests that considering the hidden heterogeneity related to farming practices in the landscape is a major factor to consider to understand natural enemy community structure than the effect of semi-natural habitats in agricultural landscapes (Vasseur et al., 2013). Recent syntheses have highlighted the strong variability in the responses of natural enemies and pests t to the proportion of semi-natural habitat with approximately the same amount of studies finding positive and negative responses (Karp et al., 2018; Tscharntke et al., 2016). We provides one key explanation for this variable effects of landscape composition by showing that farming practices within the landscape matrix are strong determinants of natural enemy meta-communities and can therefore modulate the effects of semi-natural habitats on natural enemies. Meta-community dynamics governed by spillover of predators between farming system types in the landscape are likely to explain these patterns (Vasseur et al., 2013; Leibold et al., 2004).

Our results clearly illustrate the strong taxonomic group-dependency in the observed effects of organic farming at multiple spatial scales which is in line with previous studies on other groups (Gabriel et al., 2010). Other recent studies have explored the effect of organic farming proportion on diversity of other trophic levels and revealed contrasted results (Henckel et al., 2015; Puech, Poggi, Baudry & Aviron, 2015; Petit et al., 2016; Rundlöf, Bengtsson & Smith, 2008; Inclan et al., 2015; Diekötter, Wamser, Wolters & Birkhofer, 2010; Djoudi et al., 2018). In these studies, the beneficial effect of the proportion of organic farming on plant diversity appears as a robust pattern while it seems to fade away with increasing trophic levels suggesting that bottom-up effects of plant diversity might be blurred by pest management or strongly depend on key functional traits of species (Wood et al., 2015). Our results about the taxonomic group-dependency in the effect of organic farming at multiple spatial scales highlight that there is no a unique strategy at a dedicated scale for conservation management but that strategies should be adapted to regional context (Gabriel et al., 2010).

Although organic farming at the local scale enhances abundance of natural enemies, increasing the area under organic farming at the landscape scale (here until 25% of land cover) might not be sufficient to contribute to biodiversity conservation. Indeed, among all the response variables tested in this study, the proportion of organic farming alone was never positively correlated with natural enemy abundance or diversity while local organic farming clearly increases natural enemy abundances. Moreover, we found that organic farming benefits to the most dominant species in vineyards (Figure S2). As we previously demonstrated that organic farming fosters pest control services in such vineyards (Muneret et al., 2019), it suggests that pest control might be provided by a restrictive number of dominant species. This is in line with a recent synthesis showing that pollination is provided by a restrictive number of abundant pollinator species in croplands leading to the conclusion that delivery of pollination services is an insufficient argument to conserve pollinators (Kleijn et al., 2015).

Contrary to our expectation, the proportion of semi-natural habitats had only one significant negative effect on harvestman abundance suggesting two non-exclusive explanations. First, movements of arthropods between vineyards and semi-natural habitats might be lower in such perennial landscapes compared to annual landscapes. Agrobiontes species can find food resources and refuges throughout the year in such a perennial crop and may accomplish their whole life-cycle within fields especially when spontaneous vegetation is present within the field (Rusch, Delbac & Thiéry, 2017). Spillover effects that are generally observed between semi-natural habitats and crops may therefore contribute to natural enemy assemblages within-field much more in annual than in perennial crops (Batáry, Báldi, Kleijn & Tscharntke, 2011). Second, it is possible that semi-natural habitats increase intra-guild predation by recruiting top predators and reducing arthropod natural enemy abundance within fields (Martin, et al. 2013; Barbaro et al., 2017). Such positive effects of semi-natural habitats may therefore blur the potential spillover of natural enemy communities between semi-natural habitats and crops.

Beyond the organic/non-organic farming system, specific farming practices such as pesticide use intensity, tillage intensity and crop productivity are key factors impacting natural enemy abundance and species richness. Our results clearly highlight that that reducing management intensity at the field scale is beneficial for natural enemy abundance and diversity (Winter et al., 2018). Field age was also a strong factor explaining natural enemy richness. Indeed, a long-term history assembly could be involved in the current assembly of a given community (Chase, 2003). Old vineyards could shelter species that have good ability for dispersal and species implemented for a long time, that are well adapted to the specific conditions of the field contrary to relatively new fields (Le Provost et al., 2017). We advocate that further studies conducted in perennial crops should better integrate field age as a covariate to explain biodiversity patterns at the landscape scale. These results therefore suggest that increasing field age and decreasing management intensity at the field scale are efficient management options to develop ecological intensification of vineyard independently of the type of farming systems.

### 4.1 Synthesis and applications

The diversity of farming practices at multiple spatial scales is an important aspect of landscape heterogeneity to explain natural enemies assemblages in agricultural landscapes. Considering farming practices within the landscape helps in making sense of the observed context-dependency of the effects of the proportion of semi-natural habitats on natural enemies and biological control services. We demonstrated that organic farming at the local scale has a strong effect on abundances of dominant species but not on species richness and that there is no unique response of biodiversity to organic farming expansion. All these results indicate that policies promoting the expansion of organic farming will benefit more to ecological intensification strategies seeking to enhance ecosystem services than to biodiversity conservation. Finally, we show that reducing pesticide use intensity, soil tillage intensity and crop productivity are efficient management options that benefit to natural enemies abundance and richness independently of the type of farming systems.

## ACKNOWLEDGMENTS

We are grateful to Emilie Vergnes, Laura Arias, Lisa Le Postec, Lionel Druelle, Pascale Roux, Benjamin Joubard, Delphine Binet, Nicolas Henon, Alexis Saintilan, Patrick Dauphin and Christophe Galkowski for their technical help. We thank the 38 grapevine growers for allowing us access to their vineyards. This research was funded by the Region Aquitaine and the Agence Française pour la Biodiversité (SOLUTION project). L.M. was financed by the SOLUTION project and the cluster of Excellence COTE.

## AUTHOR CONTRIBUTIONS

L.M., D.T. and A.R. conceived and designed the study. L.M., A.A., O.B., S.R.C., and A.R. collected the data. L.M. and A.R. analysed the data. L.M. wrote the first draft of the manuscript and all authors contributed substantially to the writing of the final manuscript and gave final approval for publication.

## DATA ACCESSIBILITY

Data available from Portail Data Inra: https://doi.org/10.15454/UCRZO7.

