## Supporting information for "Organic farming expansion drives natural enemy abundance not diversity in agricultural landscapes"

**FIGURE S1** Probability of each scale to explain the response variables. For each response variable, we recalculated the Akaike weights among all of the models from the four different sets ('M0', 'M1' and both 'M2' obtained using multimodel inference). We, therefore, estimated the relative importance of each level of complexity for a given response variable that gave us the "best scale" of response (figure S2). The sum of the Akaike weights ("Sum Wi") of the models obtained at the best scale provided the modelôs probability of being the top model across all of the scales.

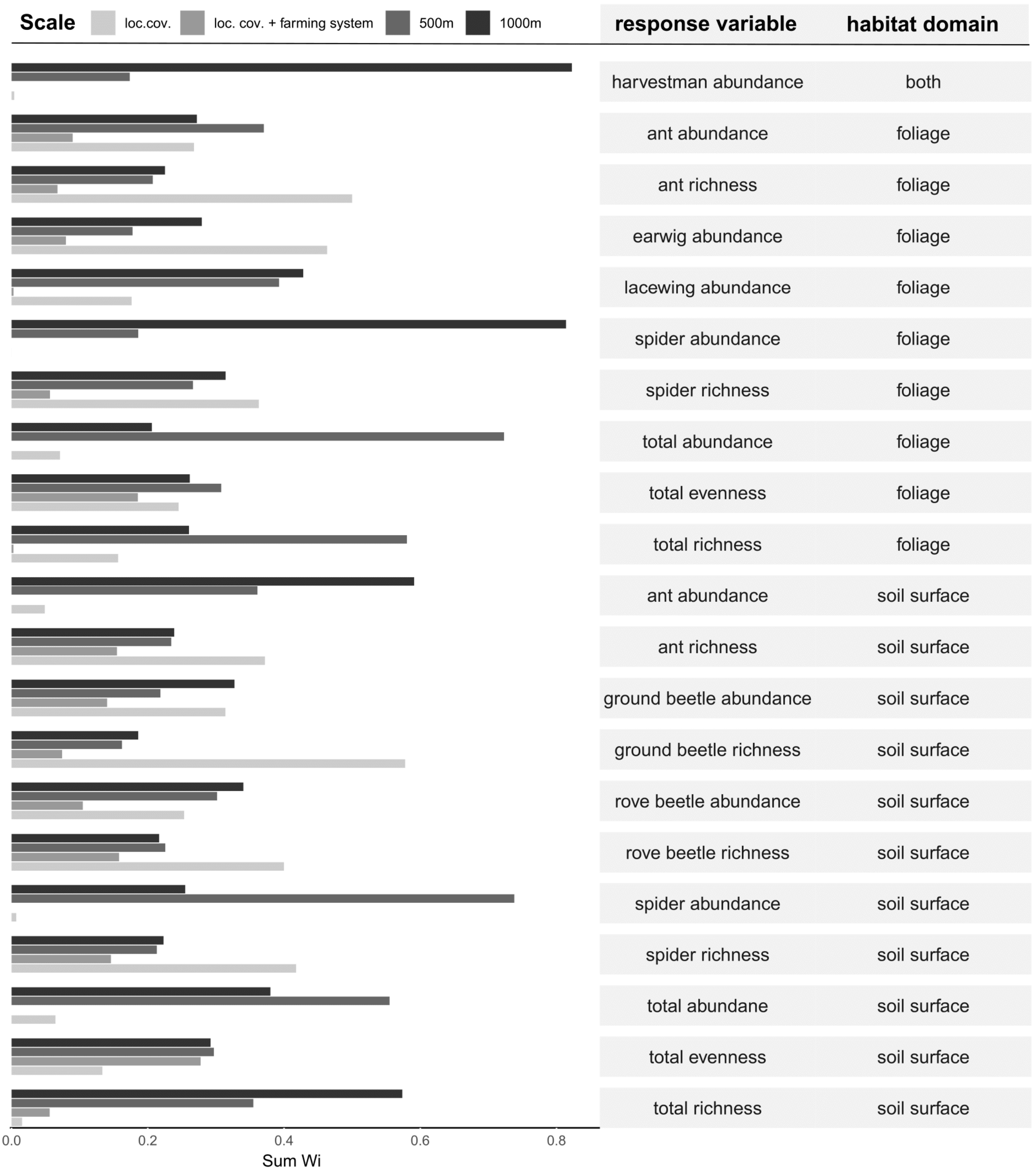

**FIGURE S2** Abundances of the 30 dominant species of the above-ground community in organic and conventional fields.

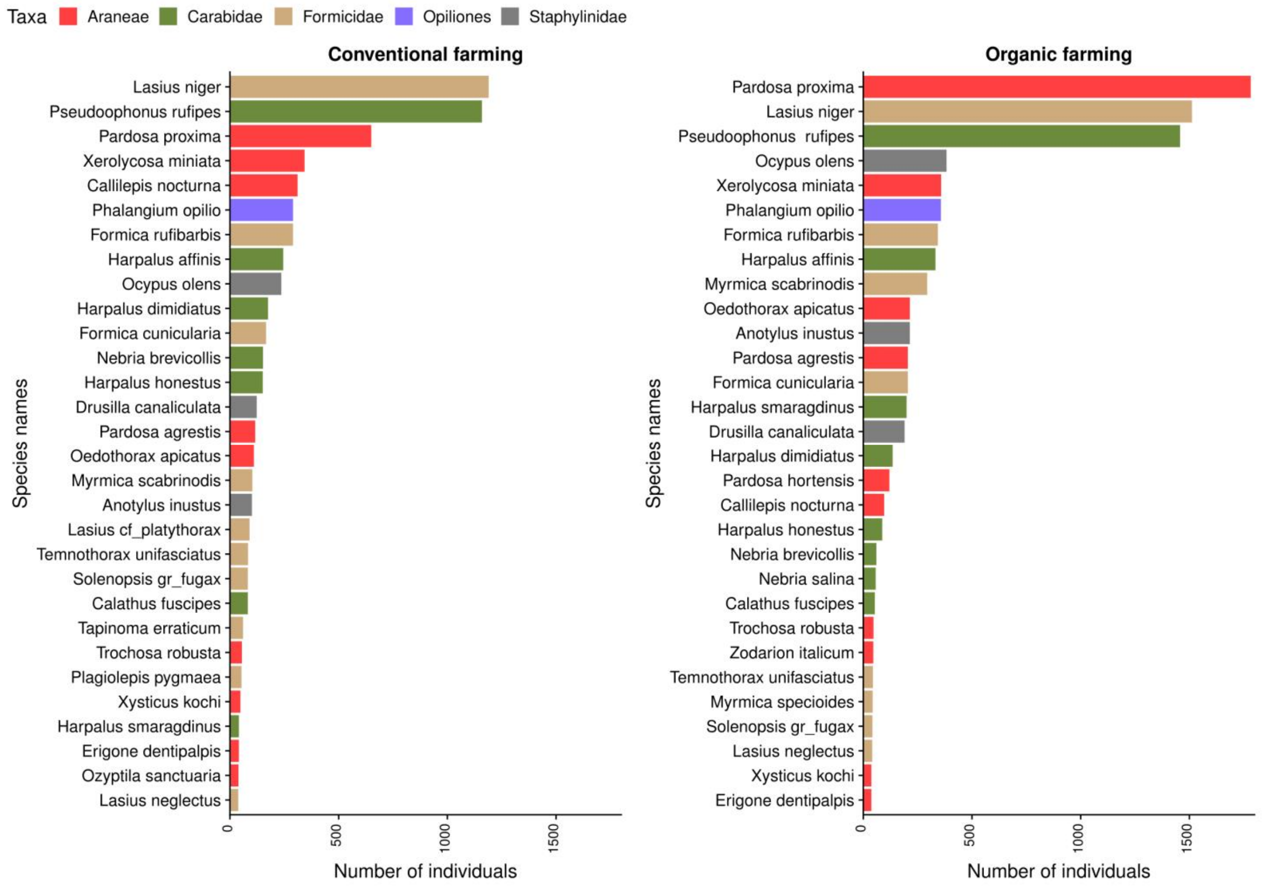
